# Kinase inhibition of G2019S-LRRK2 restores autolysosome formation and function to reduce endogenous alpha-synuclein intracellular inclusions

**DOI:** 10.1101/707273

**Authors:** Julia Obergasteiger, Giulia Frapporti, Giulia Lamonaca, Sara Pizzi, Anne Picard, Francesca Pischedda, Giovanni Piccoli, Evy Lobbestael, Veerle Baekelandt, Andrew A. Hicks, Corrado Corti, Peter P. Pramstaller, Mattia Volta

## Abstract

The Parkinson’s disease (PD)-associated kinase Leucine-Rich Repeat Kinase 2 (LRRK2) is a potent modulator of autophagy and impacts on lysosome biology and function, but unclarity exists on the precise mechanics of its role and the direction of this modulation. LRRK2 is also involved in the degradation of pathological alpha-synuclein, with pathogenic mutations precipitating neuropathology in cellular and animal models of PD, and most LRRK2 familial cases manifesting with Lewy neuropathology. Defects in autophagic processing and lysosomal degradation of alpha-synuclein have been postulated to underlie its accumulation and onset of neuropathology. Thus, it is critical to reconcile these independent pieces of information to obtain a comprehensive knowledge on LRRK2-associated pathology that could also be generalized to the idiopathic disease.

Here, we report a focused investigation on the role of PD-causing G2019S-LRRK2 in the autophagy-lysosome pathway in a recombinant cell line model. Initially, we evaluated the effect of LRRK2 expression on autophagy-related transcriptome. Then, we found that G2019S-LRRK2 leads to accumulation of autophagosomes with no net effect on autophagy induction. This is linked to abnormalities in lysosome morphology and proteolytic activity that are associated with a decrease in the successful formation of autolysosomes. Despite some of these features being shared by WT-LRRK2, alpha-synuclein intracellular inclusions are specifically found in G2019S-LRRK2 cells. Pharmacological kinase inhibition is capable of rescuing defects in the autophagy-lysosome pathway and reducing the number of inclusions. Notably, this effect is prevented by upstream blockade of autophagosome-lysosome fusion events, highlighting this step of the process as critical for alpha-synuclein clearance.

## Introduction

Mutations in the *Lrrk2* gene cause late-onset, familial Parkinson’s disease (PD) with variable penetrance that is clinically indistinguishable from idiopathic PD (iPD) but with pleomorphic pathology^1^. However, the G2019S mutation is singularly the most common genetic cause of PD worldwide, with an incidence up to 40% in specific populations^2,3^ and mostly presents with alpha-synuclein (aSyn) Lewy neuropathology at autopsy^4^.

The gene codes for Leucine-Rich Repeat Kinase 2 (LRRK2), a large multidomain protein with two distinct enzymatic domains (GTPase and kinase) in close vicinity to each other^5^. The PD-linked mutations reside in this enzymatic core with G2019S located in the kinase domain and reported to increase kinase activity^6^.

The cellular roles impacted by LRRK2 are varied, with stronger consensus on synaptic transmission^7^, vesicle trafficking^8^ and autophagy^9^. Interestingly, these pathways might converge in neuronal biology and function^10^.

Several independent investigations demonstrated that LRRK2 acts at different stages of the autophagy-lysosome pathway, with some conflicting results on the direction of this modulation^9^. Indications include a kinase-dependent role of LRRK2 in the modulation of basal autophagy, with studies showing either enhancement or repression^11–14^, and modulation of lysosome function^15,16^. In this complicated landscape, attempting to understand the impact of PD-linked mutations has further increased the complexity of the problem. Most studies indicate that mutant LRRK2 produces an aberrant autophagic function, including impairment of chaperone-mediated autophagy (CMA) and processing of aSyn^17,18^. However, macroautophagy has also been reported to be altered leading to detrimental cellular consequences^19–21^. Moreover, pathogenic LRRK2 also directly affects lysosome biology in different cell types, causing reduced lysosome function^22–25^. Thus, despite an agreement on LRRK2 playing a role in the autophagy-lysosome pathway, and that PD-linked mutations cause alteration in this process, no evidence to date indicates the precise mechanisms underlying these functions and dysfunctions.

LRRK2 has also been demonstrated to mediate accumulation of pathologic aSyn^26,27^ with kinase inhibitors being beneficial against neuropathology^26^. At the same time, aSyn neuropathology has been hypothesized to be a consequence of autophagy dysfunction (reviewed in^28^).

A missing link exists in the attempts to put these pieces together and, to the best of our knowledge, no evidence has been reported indicating how PD mutant LRRK2 specifically affects the autophagy-lysosome pathway and the consequences on aSyn handling.

Here, we set out to investigate the autophagy step(s) that are targeted by G2019S-LRRK2 and their dependence on kinase activity. We found that mutant LRRK2 alters the correct processing of autophagosomes and lysosomal activity by impairing the formation of autolysosomes. These defects are paralleled by the accumulation of endogenous aSyn in intracellular inclusions and pharmacological kinase inhibition relieves these defects. Lastly, we demonstrate that the efficacy of LRRK2 inhibition in reducing pathologic aSyn depends on the functional fusion between autophagosomes and lysosomes, indicating that this precise step is responsible for aSyn accumulation.

## Materials and methods

### Cell culture, transfection and drug treatment

SH-SY5Y neuroblastoma cell lines stably overexpressing wild-type (WT) or G2019S-LRRK2 were previously described (herein referred to as WT-LRRK2 cells and G2019S-LRRK2 cells)^29,30^. Control SH-SY5Y cells were maintained in DMEM GlutaMAX medium, supplemented with 10% fetal bovine serum (FBS) and 1% penicillin/streptomycin, while recombinant LRRK2 cells were cultured in DMEM GlutaMAX with 15% FBS, 1% non-essential amino acids, 50 μg/ml gentamycin (Gibco) and 200 μg/ml hygromycin B (Invitrogen). All cells were incubated at 37°C with 5% CO_2_.

Transfection of the GFP-LC3-mCherry reporter construct was carried out using FuGene HD (Promega) with 800 ng of DNA (ratio 5.2:1) when 50%-60% confluent, then incubated at 37°C/5% CO_2_ for 48h.

The LRRK2 kinase inhibitor PF-06447475 (herein, PF-475)^31^ was dissolved in DMSO and applied to cultured cells for 2h or 6h, while 0.1% DMSO was used as vehicle control. Chloroquine (CQ; 100 μM, 3h) was used to block to the fusion of autophagosomes with lysosomes and to evaluate the autophagic flux in cells^32^.

### Autophagy gene expression array

SH-SY5Y, WT- and G2019S-LRRK2 cells were lysed in RLT Plus buffer containing 1% β-mercaptoethanol. Total RNA was extracted using the RNeasy Plus Mini Kit (Qiagen). The RNA concentration was quantitated with QuantiFluor^®^ RNA System (Promega). First strand cDNA was synthesized from 500ng total RNA using the RT2 First Strand Kit (Qiagen) according to the manufacturers protocol. cDNA samples were then processed as indicated by the manufacturer; briefly, cDNA was mixed with RT2 SYBR^®^ Green qPCR Mastermix (Qiagen) and then dispensed to each well of the RT2 Profiler PCR Array (PAHS-084Z). Quantitative PCR was performed in a CFX96 Touch™ Real-Time PCR Detection System (BioRad). Analysis was carried out with the webtool provided by Qiagen (https://www.qiagen.com/it/shop/genes-and-pathways/data-analysis-center-overview-page/). An Excel file with CT values was uploaded to the website. Samples were assigned to control and test groups and CT values were normalized based on the geometric mean of five housekeeping genes (*ACTB, B2M, GAPDH, HPRT1* and *RPLP0*). The webtool calculates fold change using the Delta Delta C_T_ method.

### Western blotting

Cells were lysed in RIPA lysis buffer (Sigma-Aldrich) containing protease inhibitors (cOmplete tablets; Roche) and phosphatase inhibitors (PhosSTOP; Roche). Lysates were sonicated for 10 sec and then centrifuged at 10000xg for 10 minutes at 4°C. Cell lysates were heated in sample buffer (LDS sample buffer, NuPAGE) containing 50 mM dithiothreitol (DTT) at 95°C for 5 min, loaded onto a 4-12% SDS-PAGE gel and then transferred onto polyvinylidene difluoride membranes (BioRad). Primary antibodies were: anti-LRRK2 1:20000 (Abcam, ab172378), anti-pS935-LRRK2 1:1000 (Abcam, ab172382), anti-pS1292-LRRK2 1:1000 (Abcam, ab206035), anti-LC3 1:1000 (Cell Signaling Technologies, 3868), anti-β-actin 1:6000 (Sigma Aldrich, A-5316). Chemiluminescence images were acquired using Chemidoc Touch (BioRad) and relative band intensity levels were calculated using ImageLab software (BioRad).

### Immunofluorescence, confocal imaging and image analyses

Cells were fixed in 4% paraformaldehyde (PFA), then permeabilized, blocked with bovine serum albumin (BSA) and incubated with primary antibodies overnight at 4°C. The following day, after washing, cells were incubated with secondary fluorescent antibodies for 2h at room temperature, washed and mounted with DAPI. Primary antibodies were: rabbit anti-LC3B 1:2000 (Molecular Probes, L10382), mouse anti-pS129-aSyn 1:2000 (Abcam, ab184674). Secondary antibodies were: Donkey anti-Rabbit Secondary Antibody, Alexa Fluor 488 (A-21206); Donkey anti-Mouse Secondary Antibody, Alexa Fluor 555 (A31570). Visualization was performed using a Leica SP8-X confocal laser scanning microscope system equipped with an oil immersion 63X objective.

Images were analysed as projected stacks with custom pipelines in CellProfiler^33^ to quantify the number of fluorescent puncta and their integrated intensity. Our pipelines are available upon request.

### Lysotracker Deep Red and DQ-Red-BSA staining

To investigate lysosome morphology, we utilized the Lysotracker Deep Red dye (Molecular Probes, L12492) following the manufacturer’s instructions. Briefly, cells were incubated with Lysotracker for 20 min, then DAPI for live imaging (Invitrogen, R37605) was added and cells were visualized live on the confocal microscope at 63X magnification, using the *resonant* function. Single stacks were reconstructed in Imaris software (BitPlane) to a 3D structure and number and diameter of lysosomes quantified by the software.

To study the proteolytic activity of lysosomes, the fluorescent DQ-Red-BSA^34^ dye (Molecular Probes, D12051) was used following manufacturer’s instructions. Briefly, cells were incubated with the dye for 120 min, then DAPI for live imaging was added and visualization performed with confocal microscopy. Projected stack images were then analysed with CellProfiler to quantify the number of DQ-Red-BSA spots (pipeline available upon request).

### Analysis of autolysosomes with GFP-LC3-mCherry

To analyse the number of autolysosomes, we transfected recombinant LRRK2 cell lines with a GFP-LC3-mCherry construct where GFP fluorescence is pH-sensitive. Thus, in the cytosol and in autophagosomes, both GFP and mCherry fluorophores are active and both green and red fluorescence are detected. Upon fusion of the autophagosome with the lysosome and formation of the autolysosome, the pH of the vesicle turns acidic and quenches the GFP signal. Thus, autolysosomes are defined as mCherry-positive spots in confocal images.

The GFP-LC3-mCherry construct is transfected in WT-, G2019S-LRRK2 and SH-SY5Y cells using FuGene HD (see above) and fixed with 4% PFA 2 days later. Images were acquired with the confocal microscope and analysed with CellProfiler, determining the turnover from autophagosomes to autolysosomes (red/green ratio) and the number of autolysosomes (red spots). Pipeline available on request.

### Statistical analyses

Statistical analyses were performed using GraphPad Prism 8. The one-way ANOVA test was used in experiments comparing 3 or more groups, followed by Bonferroni’s *post-hoc* test for pairwise comparisons. When comparing 2 experimental groups, the unpaired, two-tailed Student’s t-test was utilized. For all analyses, the threshold for significance was set at p<0.05. All experiments were performed in a minimum of 4 independent biological replicates.

## Results

### LRRK2 alters autophagy-related transcriptome

We initially set out to evaluate general alterations to the autophagy-lysosome system in WT- and G2019S-LRRK2 cells using an unbiased approach based on transcriptome analysis of genes related to this pathway. We screened the mRNA levels of 84 genes coding for proteins playing a role at different levels in autophagy induction and initiation, vesicle formation and management, lysosome fusion and activity.

We collected the results in a clustergram and analysed general differences in expression (Fig. 1). First, we noticed that WT-LRRK2 cells mostly show opposite expression patterns with respect to G2019S-LRRK2 cells, indicating a mutation-specific effect on the transcriptome. In addition, we observed that G2019S-LRRK2 cells display differential expression of genes related to lysosome biology (e.g. *CTSD, CTSS*) and initiation of autophagy (e.g. *WIPI1, MTOR, AMBRA1, ULK1*) (Fig. 1).

**Fig. 1.**
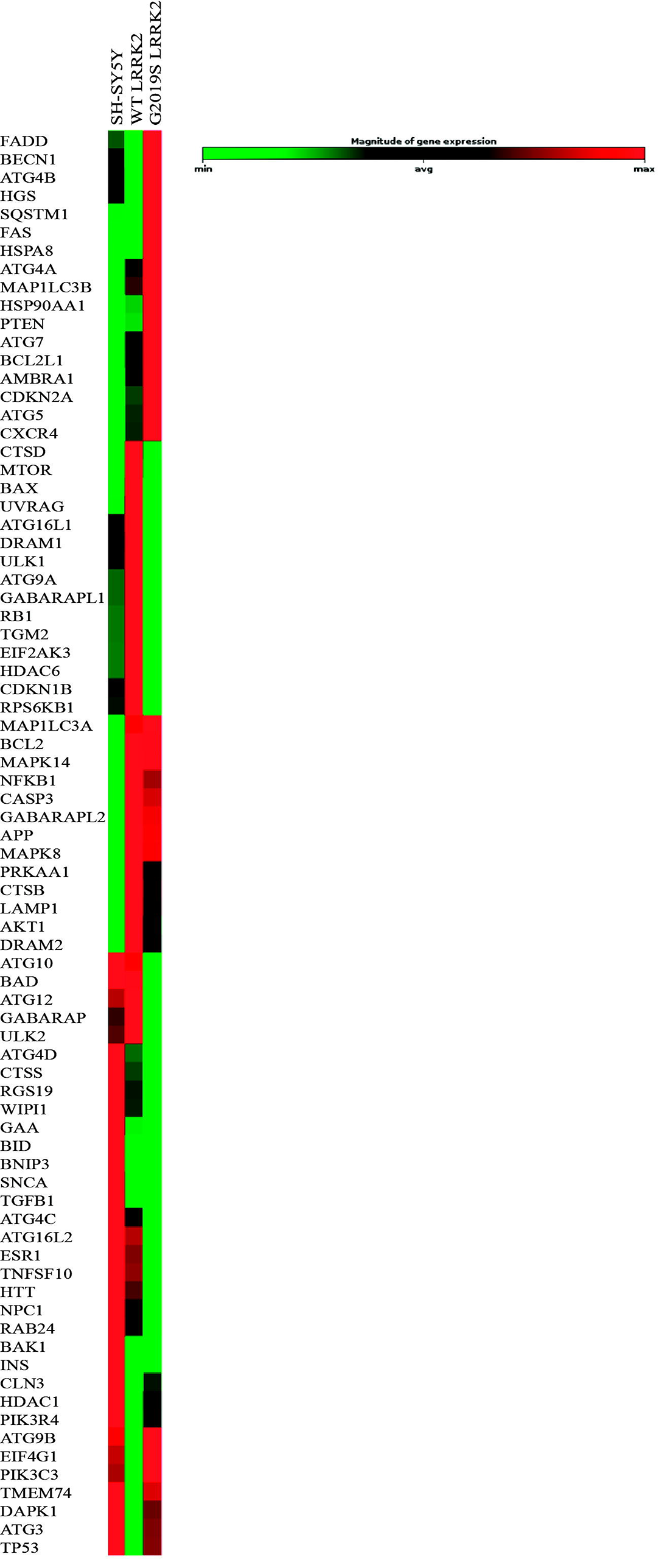
Autophagy gene expression array in LRRK2 cells. The mRNA levels of 84 autophagy-related genes were assessed via qPCR in SH-SY5Y, WT-LRRK2 and G2019S-LRRK2 cells. Of these, 78 displayed measurable expression in our cell lines and are represented as a clustergram indicating reduction (green) and increase (red) in expression. The relative difference is expressed as fold change to SH-SY5Y control cells.

Of all these genes, *WIPI1* mRNA levels have been shown to correlate directly to autophagy function and are an indicator of autophagosome formation,^35^. Then, we quantified the fold change in gene expression of WT- and G2019S-LRRK2 cells compared to SH-SY5Y controls. *WIPI1* gene expression is downregulated in WT- and G2019S-LRRK2 cells, with a stronger effect in the latter. Concomitantly, *MAP1LC3B* is upregulated (Table 1). We conclude that LRRK2 overexpression in cells affects autophagy, and the changes in gene expression might be a direct effect of LRRK2 or part of a compensatory process. The array data also suggest that the G2019S mutation operates a differential modulation and might impair the initiation of autophagy.

**Table 1.** Relative expression changes of genes related to the autophagy-lysosome pathway in WT- and G2019S-LRRK2 cells, compared to SH-SY5Y controls.

| Symbol | Fold Change (comparing to SH-SY5Y) |  |
| --- | --- | --- |
|  | WT LRRK2 | G2019S LRRK2 |
|  | Fold Change | Fold Change |
| AKT1 | 1.4747 | 1.1772 |
| AMBRA1 | 1.076 | 1.1866 |
| APP | 1.1437 | 1.1345 |
| ATG10 | 0.9823 | 0.7218 |
| ATG12 | 1.015 | 0.9341 |
| ATG16L1 | 1.2251 | 0.6987 |
| ATG16L2 | 0.9191 | 0.5776 |
| ATG3 | 0.8234 | 0.9559 |
| ATG4A | 1.0241 | 1.0386 |
| ATG4B | 0.9183 | 1.1075 |
| ATG4C | 0.7711 | 0.517 |
| ATG4D | 0.871 | 0.8158 |
| ATG5 | 1.0617 | 1.1778 |
| ATG7 | 1.3872 | 1.7698 |
| ATG9A | 1.2125 | 0.9067 |
| ATG9B | 0.7768 | 1.0211 |
| BAD | 0.9962 | 0.7481 |
| BAK1 | 0.6167 | 0.6609 |
| BAX | 1.1471 | 0.9912 |
| BCL2 | 1.721 | 1.7125 |
| BCL2L1 | 1.1551 | 1.3189 |
| BECN1 | 0.9207 | 1.1203 |
| BID | 0.762 | 0.7585 |
| BNIP3 | 0.8081 | 0.8059 |
| CASP3 | 1.8135 | 1.6912 |
| CDKN1B | 1.2985 | 0.8137 |
| CDKN2A | 3.7052 | 9.1833 |
| CLN3 | 0.8196 | 0.8844 |
| CTSB | 2.7017 | 2.0824 |
| CTSD | 1.2013 | 1.0232 |
| CTSS | 0.4744 | 0.2127 |
| CXCR4 | 7.6986 | 20.059 |
| DAPK1 | 0.8166 | 0.9504 |
| DRAM1 | 1.3401 | 0.6744 |
| DRAM2 | 1.1283 | 1.061 |
| EIF2AK3 | 1.4516 | 0.8169 |
| EIF4G1 | 0.886 | 1.0238 |
| ESR1 | 0.8725 | 0.4806 |
| FADD | 0.9149 | 1.1835 |
| FAS | 0.9578 | 1.8232 |
| GAA | 0.8453 | 0.8091 |
| **GABARAP** | 1.0485 | 0.8966 |
| **GABARAPL1** | 1.3248 | 0.8569 |
| **GABARAPL2** | 1.187 | 1.1665 |
| **HDAC1** | 0.8974 | 0.9478 |
| **HDAC6** | 1.336 | 0.8641 |
| **HGS** | 0.8673 | 1.1755 |
| **HSP90AA1** | 1.1748 | 1.7913 |
| **HSPA8** | 0.9984 | 1.7976 |
| **HTT** | 0.912 | 0.7051 |
| **INS** | 0.2576 | 0.3606 |
| **LAMP1** | 1.1317 | 1.0807 |
| **MAP1LC3A** | 1.2958 | 1.3162 |
| **MAP1LC3B** | 1.3364 | 1.5 |
| **MAPK14** | 1.3022 | 1.2991 |
| **MAPK8** | 1.5531 | 1.5069 |
| **MTOR** | 1.0465 | 1.0032 |
| **NFKB1** | 1.0485 | 1.0382 |
| **NPC1** | 0.938 | 0.8411 |
| **PIK3C3** | 0.7689 | 1.059 |
| **PIK3R4** | 0.8715 | 0.9327 |
| **PRKAA1** | 1.2937 | 1.164 |
| **PTEN** | 1.0761 | 1.3675 |
| **RAB24** | 0.939 | 0.8338 |
| **RB1** | 1.1194 | 0.9508 |
| **RGS19** | 0.8857 | 0.8221 |
| **RPS6KB1** | 1.0645 | 0.963 |
| **SNCA** | 0.7683 | 0.7474 |
| **SQSTM1** | 0.9791 | 1.0981 |
| **TGFB1** | 0.6876 | 0.6546 |
| **TGM2** | 2.0779 | 0.5568 |
| **TMEM74** | 0.6818 | 0.9542 |
| **TNFSF10** | 0.8986 | 0.5604 |
| **TP53** | 0.7308 | 0.9354 |
| **ULK1** | 1.0795 | 0.9113 |
| **ULK2** | 1.093 | 0.7751 |
| **UVRAG** | 1.1362 | 0.972 |
| **WIPI1** | 0.5497 | 0.3027 |

### Autophagy-lysosome alterations in G2019S-LRRK2 cells

The transcriptome analysis indicated probable alterations in the autophagy-lysosome pathway in our cell lines. However, autophagy is a highly dynamic process and, while transcriptomics is indicative of changes, further functional evidence is needed. Thus, to investigate whether autophagy and/or lysosome biology are functionally affected in G2019S-LRRK2 cells, we assessed autophagic flux by Western blotting (WB) for LC3B upon treatment with CQ or vehicle (Fig. 2A). In vehicle-treated cells, levels of LC3B-II are increased in G2019S-LRRK2 cells, with respect to SH-SY5Y controls (Fig. 2B). This indicates an increased number of autophagosomes. Upon treatment with CQ, formation of autolysosomes is inhibited and the conversion from LC3B-I to LC3B-II provides information on the autophagic flux^36^. The ratio of LC3B-II/LC3B-I was not different in our LRRK2 cell lines treated with CQ (Fig. 2C).

**Fig. 2.**
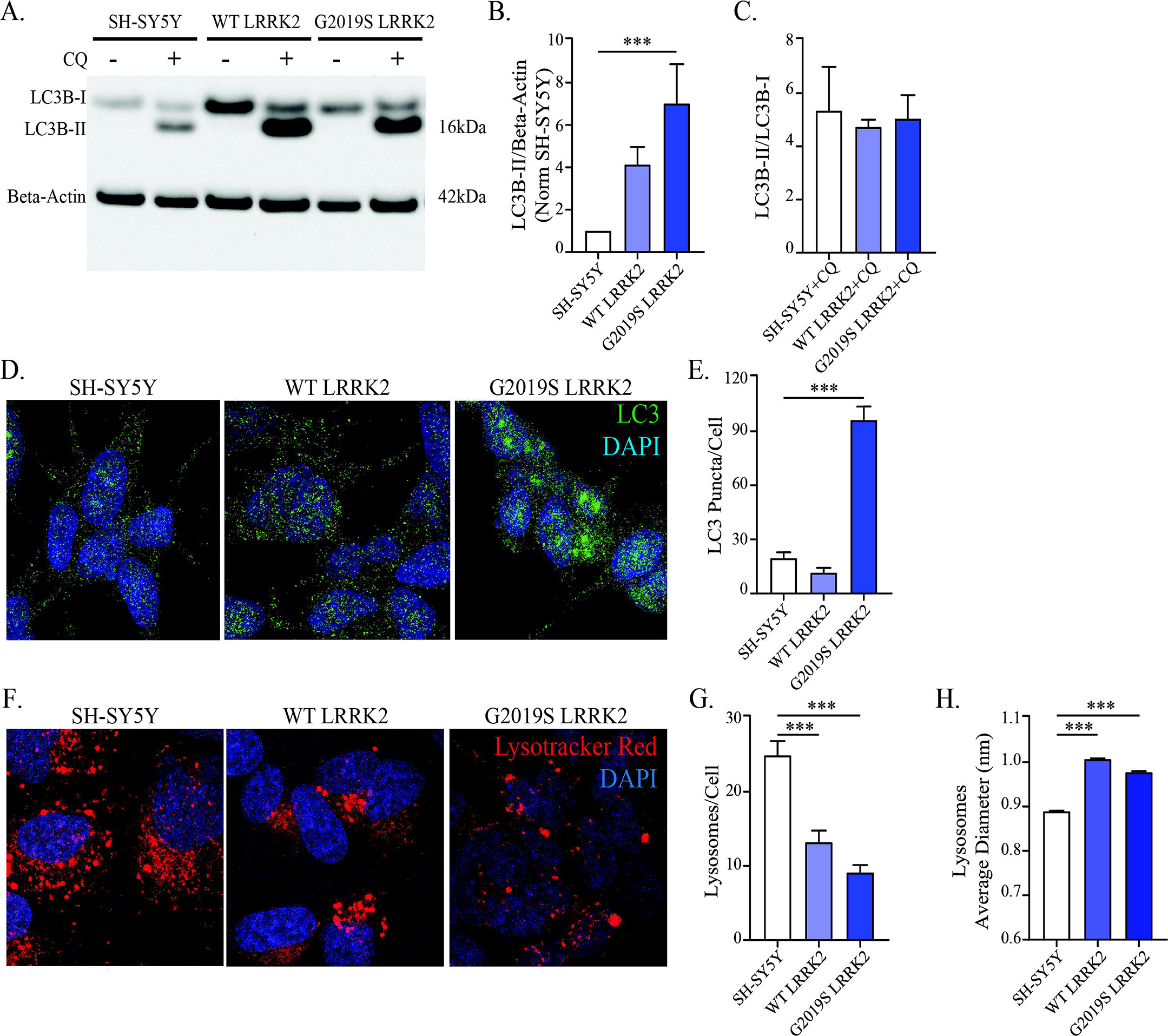
Alterations in the autophagy-lysosome pathway in recombinant LRRK2 neuroblastoma cells. A) The autophagic flux was assessed in WT- and G2019S-LRRK2 cells upon treatment with CQ (100 μM, 3h) and western blotting for LC3B. B) Quantification of LC3B levels (normalized to β-actin) reveled an increase in untreated G2019S-LRRK2 cells, indicative of an increased number of autophagosomes. C) The ratio between LC3B-II and LC3B-I was not different across CQ-treated cells, suggesting no differences in autophagic flux. D) Immunocytochemistry for LC3B was performed to visualize its endogenous distribution and autophagosomes in WT- and G2019S-LRRK2 cells. E) Quantification of LC3B-positive puncta revealed a significant increase in G2019S-LRRK2 cells. F) Processing of cells with the Lysotracker Red dye was performed to visualize lysosomes in WT- and G2019S-LRRK2 cells. G) The number of lysosomes per cell was quantified and revealed a decrease in both WT- and G2019S-LRRK2 cell lines. H) The average diameter of lysosomes was assessed in parallel. A significant enlargement was observed in both WT- and G2019S-LRRK2 cells. Data are means±SEM of 4-5 independent experiments for WB. In imaging experiments, 4-5 independent biological replicates were performed and analysis conducted on 700-1000 cells per group in each experiment. ***p<0.001, one-way ANOVA followed by Bonferroni’s *post-hoc* test.

To further investigate autophagic mechanisms, we turned to immunocytochemistry (ICC), stained for endogenous LC3B and counted the number of puncta (Fig. 2D). No difference was detected in cells overexpressing WT-LRRK2, while a strong increase in the number of LC3B puncta per cell was observed in G2019S-LRRK2 cells (Fig. 2E).

These data indicate that autophagosomes accumulate in G2019S-LRRK2 cells, but this is likely not caused by an increase in their production. To study the opposite end of the process, we used Lysotracker Red staining to visualize lysosomes (Fig. 2F). Both WT- and G2019S-LRRK2 cells displayed a significant reduction in the number of lysosomes (Fig. 2G) and a concomitant increase in their size (Fig. 2H).

Thus, we hypothesized that LRRK2 could affect the efficacy of the fusion process between autophagosomes and lysosomes, and the consequent production of autolysosomes. To assess the capacity of the cells to correctly produce autolysosomes, we transfected a double-tagged GFP-LC3-mCherry construct where the GFP fluorescence is quenched in acidic pH (see *Materials and Methods*). Using confocal microscopy, we imaged transfected cells and measured the ratio between GFP and mCherry puncta and the number of autolysosomes (mCherry puncta per cell), in order to obtain information on the processing of the exogenous LC3 (Fig. 3). Quantitative determinations indicated a significant increase of the mCherry/GFP ratio in WT-LRRK2 cells, while the G2019S mutation abolished this effect (Fig. 3B). Consistently, WT-LRRK2 cells displayed a strong increase in autolysosome formation, that is completely prevented by the G2019S mutation (Fig. 3C). These results indicate that LRRK2 overexpression impacts on the autophagic-lysosome pathway by acting on the fusion of autophagosomes with lysosomes. Altogether, our data indicate that overexpression of G2019S-LRRK2 leads to accumulation of autophagosomes without an induction of autophagy. Concomitantly, lysosomal depletion and morphological abnormalities are detected, in addition to incorrect processing of the pathway as indicated by reduced number of autolysosomes. These data suggest lysosome biology and the fusion process are affected by LRRK2. Despite WT-LRRK2 sharing some effects on lysosomal biology, G2019S-LRRK2 cells specifically accumulate undegraded autophagosomes and lose the ability to promote autolysosome formation.

**Fig. 3.**
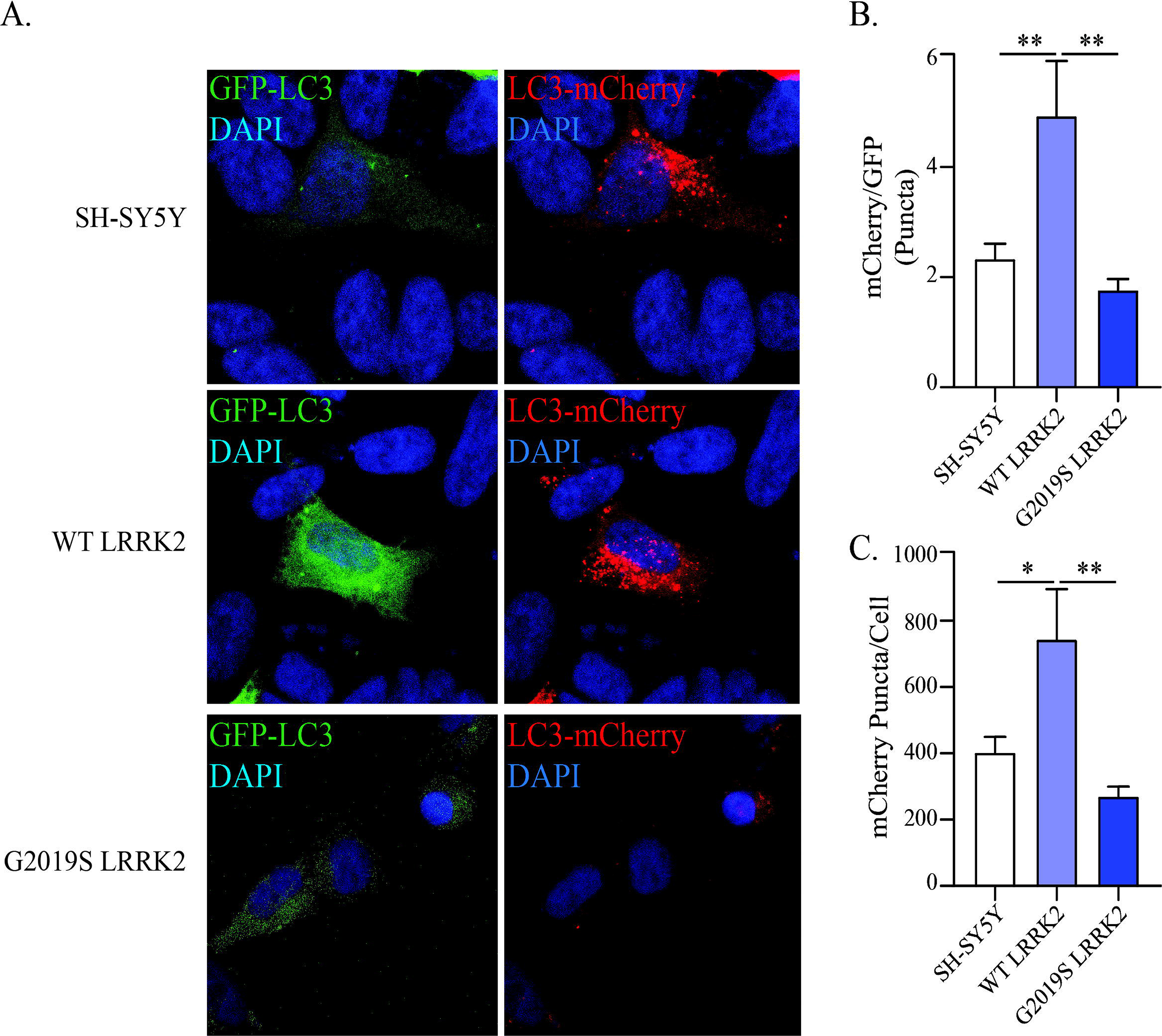
G2019S-LRRK2 cells display impaired autolysosome formation. A) The GFP-LC3-mCherry reporter construct was transfected in WT-, G2019S-LRRK2 and SH-SY5Y control cells and visualized under the confocal microscope to examine the abundance of autophagosomes and autolysosomes. B) The ratio between mCherry and GFP puncta count was determined and revealed a significant increase in fusion in WT-LRRK2 cells, which was lost in G2019S-LRRK2. C) The number of mCherry-positive autolysosomes per cell was quantified and paralleled the mCherry/GFP ratio. WT-LRRK2 cells displayed significantly more autolysosomes, while this increase was prevented by the G2019S mutation. Data are means±SEM of 4-5 independent experiments and analysis conducted on ~100 cells per group in each experiment. *p<0.05, **p<0.01, ***p<0.001, one-way ANOVA followed by Bonferroni’s *post-hoc* test.

### G2019S-LRRK2 alters lysosomal functionality to promote accumulation of intracellular alpha-synuclein inclusions

The alterations in lysosomal morphology and ability to correctly form autolysosomes observed in G2019S-LRRK2 cells prompted us to hypothesize the function of lysosomes could be diminished as a consequence. To study proteolytic activity, we employed the DQ-Red-BSA assay^34^. This protein complex carries several fluorophores that auto-quench each other. The DQ-Red-BSA is endocytosed and trafficked to lysosome where the hydrolases degrade it and the fluorophores become isolated and thus fluorescent. Thus, enhancement of the fluorescence signal indicates stronger proteolytic activity. After incubation with the substrate, living cells were imaged with a confocal microscope (Fig. 4A). Both WT- and G2019S-LRRK2 cells displayed an increase in the number of DQ-Red-BSA puncta, with respect to SH-SY5Y controls (Fig. 4B) indicating that high LRRK2 levels enhance the proteolytic activity of lysosomes. However, G2019S-LRRK2 cells had significantly fewer DQ-Red-BSA spots with respect to WT-LRRK2. We conclude that the G2019S mutation produces a defect in lysosomal activity, when compared to WT-LRRK2. These results are consistent with what observed for autolysosome formation (Fig. 3).

**Fig. 4.**
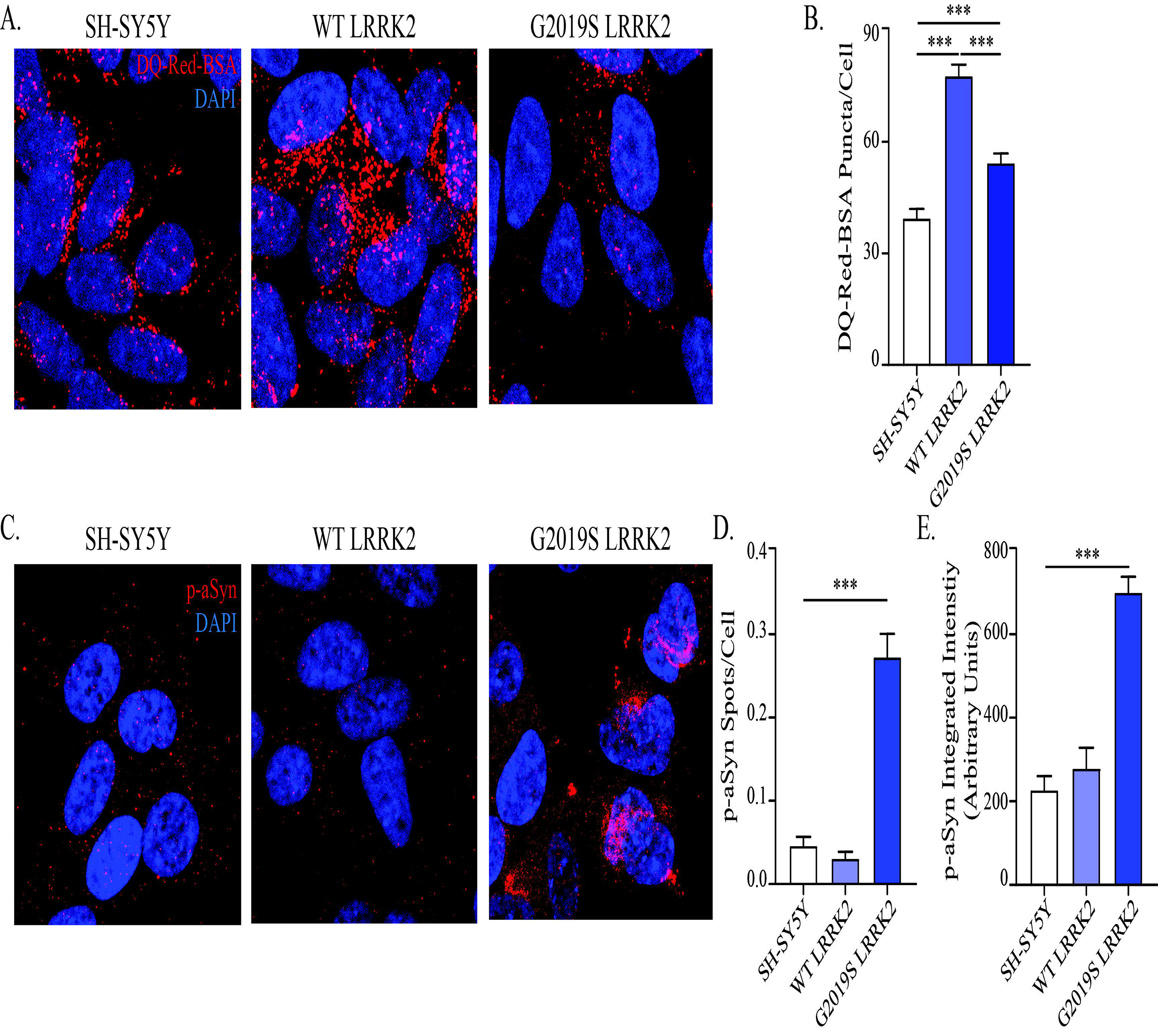
G2019S-LRRK2 cells display impaired lysosome activity and accumulate intracellular pS129-aSyn inclusions. A) The DQ-Red-BSA assay was employed to assess the proteolytic activity of lysosomes in WT- and G2019S-LRRK2 cells. B) Quantification of DQ-Red-BSA fluorescent spots revealed a significant increase in WT-LRRK2 cells, while G2019S-LRRK2 cells displayed significantly less spots when compared to WT-LRRK2 cells. C) Immunocytochemistry for pS129-aSyn was employed to investigate the presence of intracellular aSyn inclusions. D) The number of inclusions was quantified and G2019S-LRRK2 cells specifically displayed discrete intracellular objects. E) The integrated intensity of the immunosignal was also measured and revealed a strong increase in G2019S-LRRK2 cells. Data are means±SEM of 4-5 independent experiments and analysis conducted on 700-1000 cells per group in each experiment. ***p<0.001, one-way ANOVA followed by Bonferroni’s *post-hoc* test.

The DQ-Red-BSA assay is informative on the general capacity of cells to degrade protein cargoes via the lysosomes, but does not provide information on specific substrates whose degradation might be affected. PD patients with G2019S mutation in LRRK2 predominantly exhibit Lewy pathology^3,4^ and in experimental models this mutation has been shown to promote accumulation of pathological alpha-synuclein^26^. Moreover, our transcriptome analysis showed altered mRNA levels of *CTSB*, encoding for the lysosomal protease Cathepsin B, which plays an essential role for aSyn degradation^37^. Thus, we probed our cell lines for the presence of pathologic pS129-aSyn (Fig. 4C). Control SH-SY5Y and WT-LRRK2 cells displayed only diffuse immunostaining for pS129-aSyn that did not resemble any organized or higher molecular structure. Conversely, stronger staining was observed in G2019S-LRRK2 cells that appeared as intracellular inclusions, mostly around the nucleus. Quantification of the number of pS129-aSyn spots per cell yielded a strong increase in G2019S-LRRK2 cells (Fig. 4D), in parallel with a significant enhancement of the integrated intensity of the immunosignal (Fig. 4E).

In summary, endogenous aSyn accumulates specifically in G2019S-LRRK2 cells, probably due to impaired lysosomal degradation following ineffective formation of autolysosomes.

### Pharmacological kinase inhibition of G2019S-LRRK2 reduces accumulation of autophagosomes and restores lysosome function

The G2019S mutation has been linked to a toxic increase in LRRK2 kinase activity^6^ and this has led to the development of LRRK2-selective kinase inhibitors as experimental therapeutic strategy for PD^38^. Thus, we sought to investigate the kinase-dependence of the phenotypes observed in our G2019S-LRRK2 cell line specifically, as the one displaying accumulation of aSyn. Of note, pharmacological kinase inhibition did not dramatically affect the autophagy-related transcriptome (Supp. Table 1). This is expected in the light of the short application time (max 6h).

First, we evaluated the level of phosphorylation of LRRK2 at the S1292 residue, which is an autophosphorylation site directly related to kinase activity^39^, in all our cell lines (Fig. 5A). Expression of endogenous LRRK2 in control SH-SY5Y cells is extremely weak and rendered quantification of optical density highly inaccurate. We observed a strong increase of active LRRK2, measured as the ratio between pS1292-LRRK2 and total LRRK2 protein, in G2019S-LRRK2 cells when compared to WT-LRRK2 cells (Fig. 5B). Interestingly, we did not observe an increase in pS935-LRRK2 (but rather a significant reduction; Supp. Fig. 2), indicating that phosphorylation at this residue is not informative on kinase activity in our hands. Next, we tested the selective LRRK2 inhibitor PF-475 for its ability to induce dephosphorylation at both S1292 and S935 residues, which are both targets of pharmacological kinase inhibition^31,39^. PF-475 (300 nM-1 μM) reduced phosphorylation at both serine residues (Fig. 5C). Given the strong inhibition of pS1292-LRRK2 in G2019S-LRRK2 cells observed at 300 nM, we decided to use 300 nM and 500 nM concentrations in subsequent experiments. This strategy limited the destabilization of LRRK2 protein observed after treatment with 1 μM PF-475, an effect we expected based on the literature^40^.

**Fig. 5.**
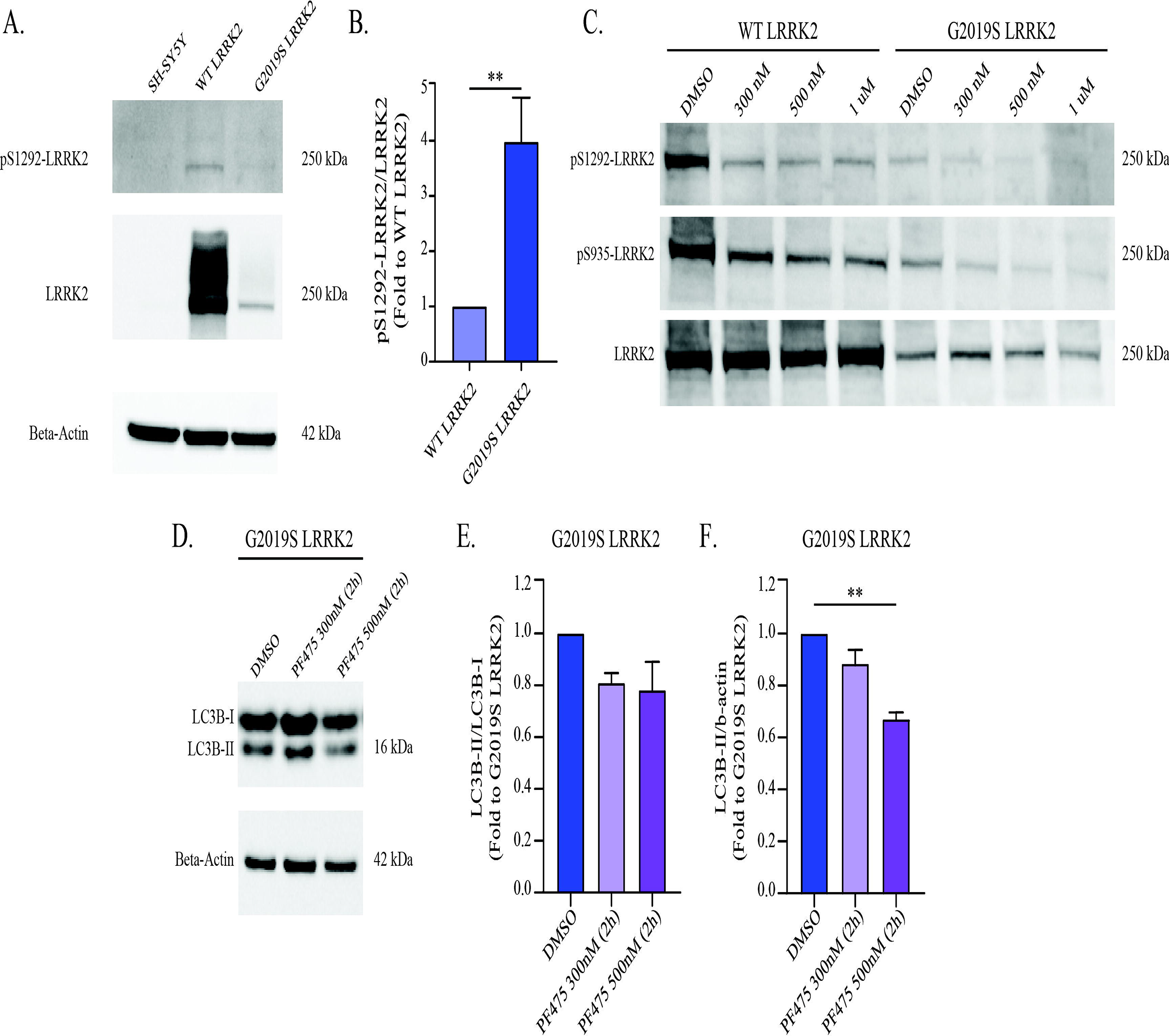
LRRK2 kinase inhibition in G2019S-LRRK2 cells does not affect autophagic flux but reduces LC3B-II levels. A) Western blot for pS1292 autophosphorylation site of LRRK2 was performed to assess LRRK2 kinase activity in our cell lines. B) The pS1292-LRRK2/LRRK2 optical density ratio indicated that LRRK2 kinase activity is strongly enhanced in G2019S-LRRK2 cells when compared to WT-LRRK2. C) WT- and G2019S-LRRK2 cells were treated with the PF-475 LRRK2 kinase inhibitor (300 nM-1 μM) for 2h, then processed for Western blot to evaluate dephosphorylation of S1292 and S935. D) G2019S-LRRK2 cells were treated with PF-475 (300-500 nM, 2h) and processed for Western blot for LC3B to evaluate the effect on autophagy initiation. E) LRRK2 kinase inhibition was not effective on the autophagic flux, as assessed by the conversion of LC3B-I to LC3B-II. F) The number of autophagosomes, as assessed by LC3B-II levels, was reduced by PF-475 500 nM in G2019S-LRRK2 cells. Data are means±SEM of 4-5 independent experiments. **p<0.01, unpaired two-tailed Student’s t-test (Panel B). **p<0.01, one-way ANOVA followed by Bonferroni’s *post-hoc* test.

We assessed the effect of LRRK2 kinase inhibition on autophagic flux in G2019S-LRRK2 cells (Fig. 5D-F). After 2h application of PF-475, cells were lysed and subjected to WB to measure LC3B levels (Fig. 5D). The conversion from LC3B-I to LC3B-II was not changed in PF-475-treated cells in our conditions (Fig. 5E), indicating the autophagic flux is not affected. Conversely, LC3B-II levels were significantly reduced by PF-475 at 500 nM (Fig. 5F), suggesting a decrease in the number of autophagosomes.

To substantiate this finding, we applied PF-475 before processing the cells for confocal microscopy. After 2h treatment, we immunostained G2019S-LRRK2 cells for LC3 and counted the number of puncta (Fig. 6A). Both 300 nM and 500 nM PF-475 significantly reduced the number of LC3-positive puncta in G2019S-LRRK2 cells (Fig. 6B). In addition, we assessed the involvement of increased kinase activity in the lysosomal defects we observed in G2019S-LRRK2 cells, and the efficacy of PF-475 in modifying those, using Lysotracker Red staining (Fig. 6C). The number of lysosomes per cell was significantly increased by the 500 nM concentration after 2h (Fig. 6D), while both 300 nM and 500 nM concentrations of PF-475 were capable of decreasing lysosomal size (Fig. 6E).

**Fig. 6.**
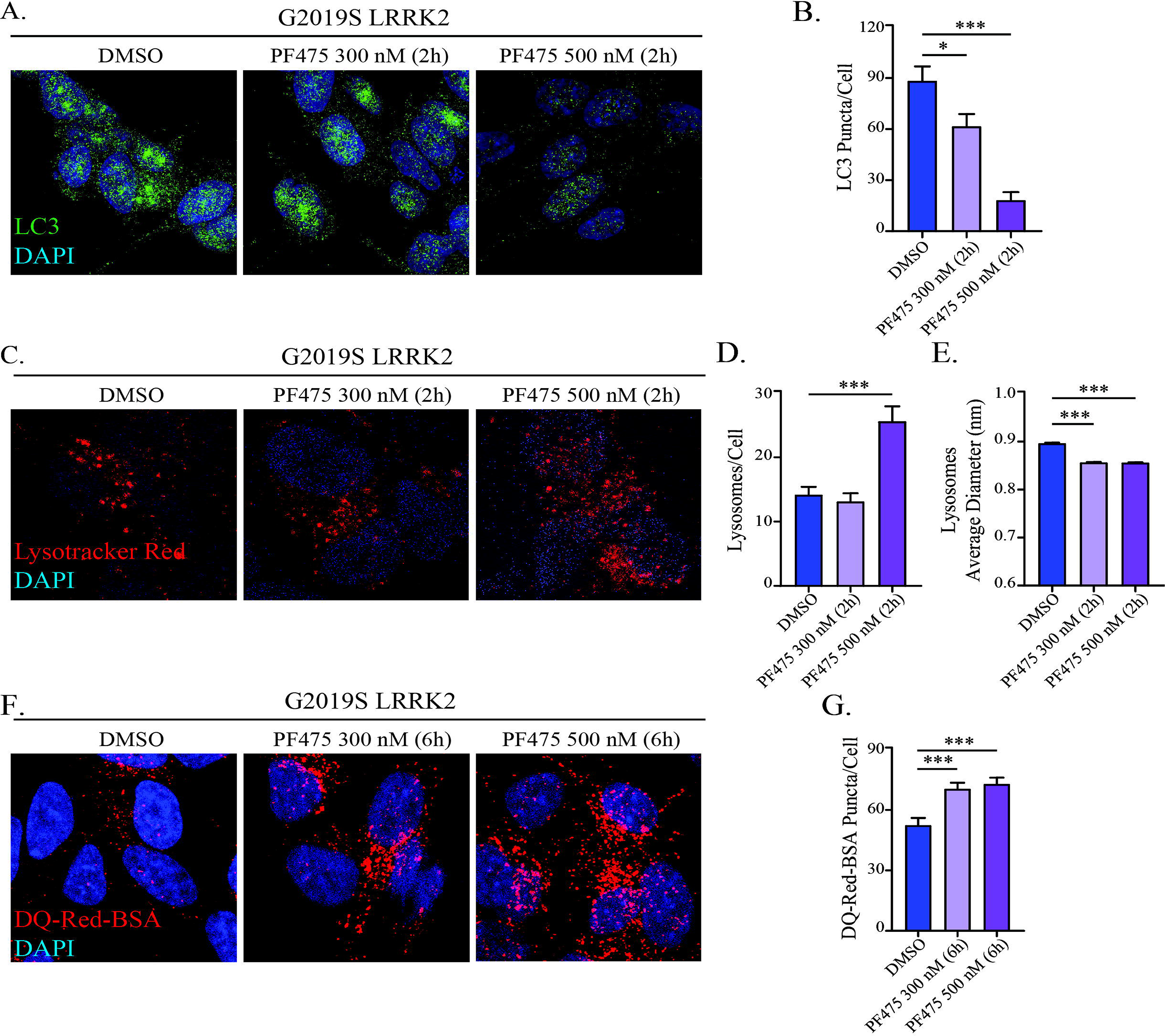
LRRK2 kinase inhibition in G2019S-LRRK2 cells ameliorates autophagosome clearance, lysosome morphology and degradative capacity. A) G2019S-LRRK2 cells were treated with PF-475 (2h) and subjected to immunocytochemistry for LC3B. B) Quantification of LC3B-positive puncta revealed that PF-475 concentration-dependently reduced the number of puncta in G2019S-LRRK2 cells. C) G2019S-LRRK2 cells treated with PF-475 were incubated with the Lysotracker Red dye to visualize lysosomes. D) Quantification of lysosome number per cell revealed that PF-475 500 nM significantly increased their abundance in G2019S-LRRK2 cells. E) Determination of lysosome diameter revealed that PF-475, at both concentrations, attenuated the enlargement observed in G2019S-LRRK2 cells. F) G2019S-LRRK2 cells were treated with PF-475 for 6h, then incubated with DQ-Red-BSA and imaged via confocal microscopy to assess proteolytic activity. G) Quantification of the number of DQ-Red-BSA spots per cell revealed that both concentrations were effective in enhancing lysosome degradative capacity in G2019S-LRRK2 cells. Data are means±SEM of 4-5 independent experiments and analysis conducted on 700-1000 cells per group in each experiment. *p<0.05, ***p<0.001, one-way ANOVA followed by Bonferroni’s *post-hoc* test.

Using DQ-Red-BSA after exposure to PF-475 we observed that LRRK2 kinase inhibition also affected lysosomal activity (Fig. 6F). For these experiments, we applied PF-475 for 6h to compensate for the necessary 2h incubation with DQ-Red-BSA. Both concentrations (300 nM and 500 nM) significantly enhanced the number of fluorescent spots per cell (Fig. 6G), indicating that kinase inhibition rescued the proteolytic deficits in G2019S-LRRK2 cells.

Collectively, these data demonstrate that inhibition of G2019S-LRRK2 kinase function rescues lysosomal abnormalities and restores the correct degradation of autophagosomes.

### LRRK2 kinase inhibition reinstates correct autolysosome formation and promotes degradation of aSyn inclusions

Having observed positive effects exerted by PF-475 on autophagosome number, lysosome morphology and functionality, we next asked what specific step of the autophagic process is modulated by LRRK2 kinase inhibition in G2019S-LRRK2 cells. For this, we used the GFP-LC3-mCherry construct to investigate whether the impairment in autolysosome formation also depends on kinase overactivity (Fig. 7A). G2019S-LRRK2 cells transfected with this reporter construct and exposed to PF-475 (300 nM-500 nM, 6h) were imaged with confocal microscopy and analyzed as reported above. The mCherry/GFP ratio was enhanced by PF-475 (Fig. 7B), indicating an increase in autophagosome processing and reinstatement of the correct fusion events. Consistently, the number of mCherry-positive autolysosomes was strongly increased by PF-475, proportionally to the concentration used but reaching statistical significance at 500 nM (Fig. 7C).

**Fig. 7.**
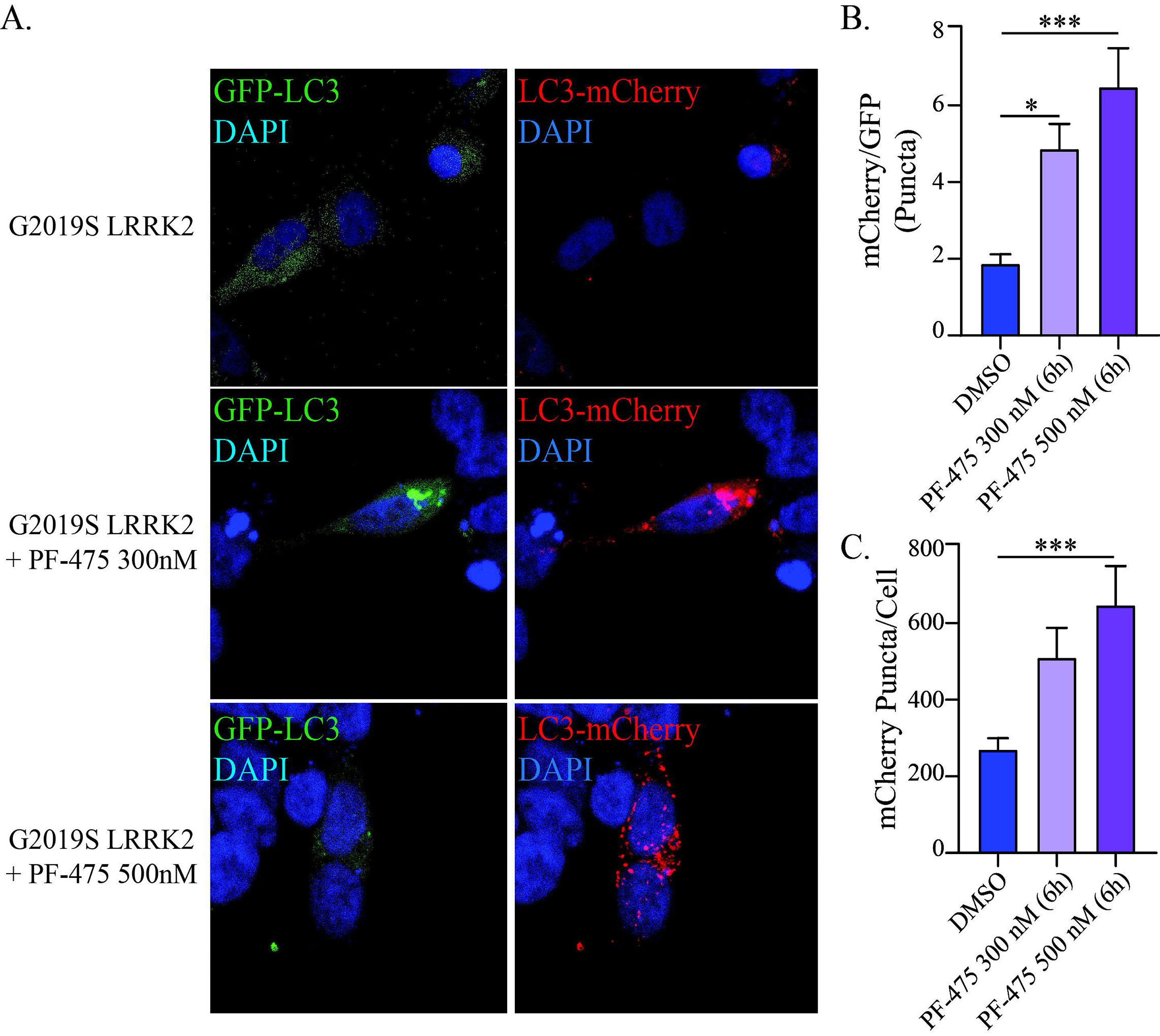
LRRK2 kinase inhibition in G2019S-LRRK2 cells promotes the formation of autolysosomes. A) G2019S-LRRK2 cells were transfected with the GFP-LC3-mCherry reporter and visualized with confocal microscopy. B) The ratio between mCherry and GFP puncta count was determined and revealed a significant increase in fusion after treatment with PF-475 at both 300 nM and 500 nM concentrations. C) The number of mCherry-positive autolysosomes per cell was quantified and showed an increase after treatment with PF-475, that reached statistical significance at 500 nM. Data are means±SEM of 4-5 independent experiments and analysis conducted on ~100 cells per group in each experiment. *p<0.05, ***p<0.001, one-way ANOVA followed by Bonferroni’s *post-hoc* test.

At this point, we sought to determine if the effects of PF-475 on lysosomal activity and the autophagic processing was also affecting aSyn inclusions in G2019S-LRRK2 cells. We treated G2019S-LRRK2 cells with PF-475 for 2h and then processed them for pS129-aSyn immunostaining (Fig. 8). In PF-475-treated cells, pS129-aSyn staining appeared mostly diffuse as opposed to larger structures identified in DMSO control (Fig. 8A). Both 300 nM and 500 nM concentrations significantly reduced the number of pS129-aSyn spots (Fig. 8B) and the integrated intensity of the fluorescent signal (Fig. 8C). These effects were maintained also after 6h treatment with PF-475 (Supp. Fig. 3), consistent with what we observed in the GFP-LC3-mCherry and DQ-Red-BSA experiments described above.

**Fig. 8.**
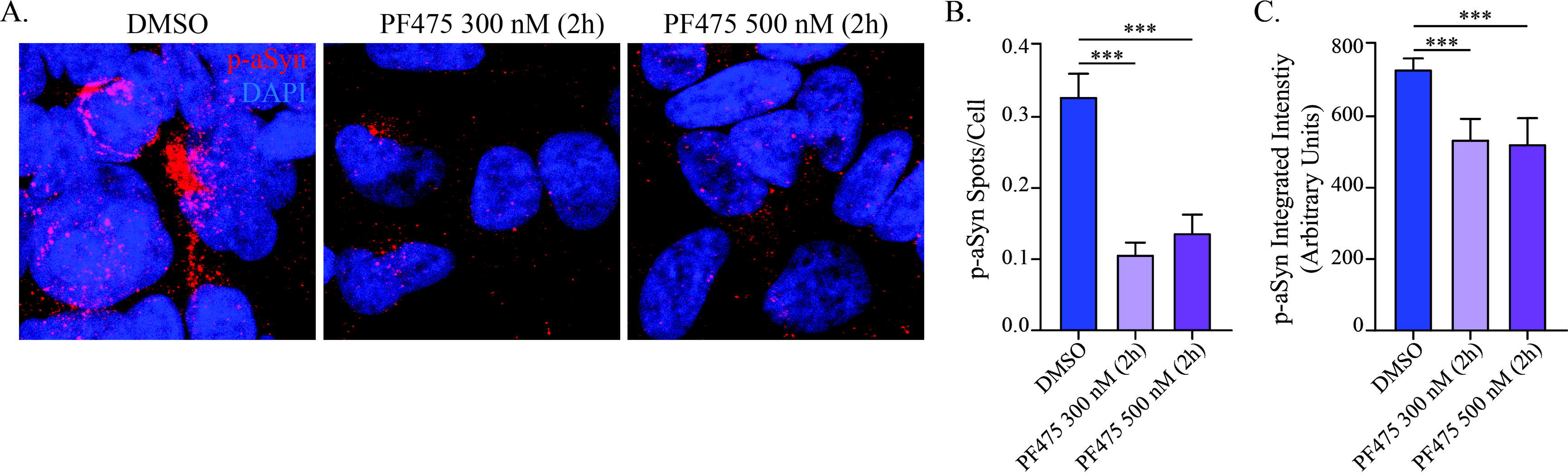
LRRK2 kinase inhibition in G2019S-LRRK2 cells reduces pS129-aSyn intracellular inclusions. A) G2019S-LRRK2 cells were treated with PF-475 for 2h and processed for immunocytochemistry for pS129-aSyn. B) The number of pS129-aSyn inclusions per cell was quantified and revealed a significant reduction operated by both 300 nM and 500 nM PF-475. C) The integrated intensity of the immunosignal was assessed at the same time and revealed both PF-475 concentrations were effective in reducing pS129-aSyn intensity in G2019S-LRRK2 cells. Data are means±SEM of 4-5 independent experiments and analysis conducted on 700-1000 cells per group in each experiment. ***p<0.001, one-way ANOVA followed by Bonferroni’s *post-hoc* test.

Therefore, inhibition of G2019S-LRRK2 kinase activity rescues autophagy-lysosome deficits and enables the cells to process aSyn, reducing its accumulation in intracellular structures, specifically through promotion of correct autophagosome-lysosome fusion.

### PF475-mediated reduction of pS129-aSyn inclusions depends on correct fusion of autophagosomes with lysosomes

Our results suggest that G2019S-LRRK2 induces abnormalities in the autophagic process with specific impairment of lysosome function and leads to accumulation of intracellular pS129-aSyn. These deficits appear to originate from an inefficient formation of autolysosomes. We then sought to determine if the effect of PF-475 on aSyn inclusions directly depends on the correct fusion of autophagosomes with lysosomes. Since CQ inhibits lysosomal activities through impairing this critical fusion step^32^, we applied CQ (100 μM, 3h) to G2019S-LRRK2 cells to block autolysosome formation prior to exposure to PF-475 (500 nM, 2h), then processed the cells for ICC and confocal microscopy (Fig. 9A). We confirmed the reduction of pS129-aSyn inclusions after application of PF-475 alone. CQ alone induced a non-significant trend towards an increased number of inclusions. Importantly, PF-475 in CQ-treated cells lost its efficacy and did not further alter the number of pS129-aSyn inclusions (Fig. 9B). The same effects were evident also when measuring the intensity of the immunosignal (Fig. 9C).

**Fig. 9.**
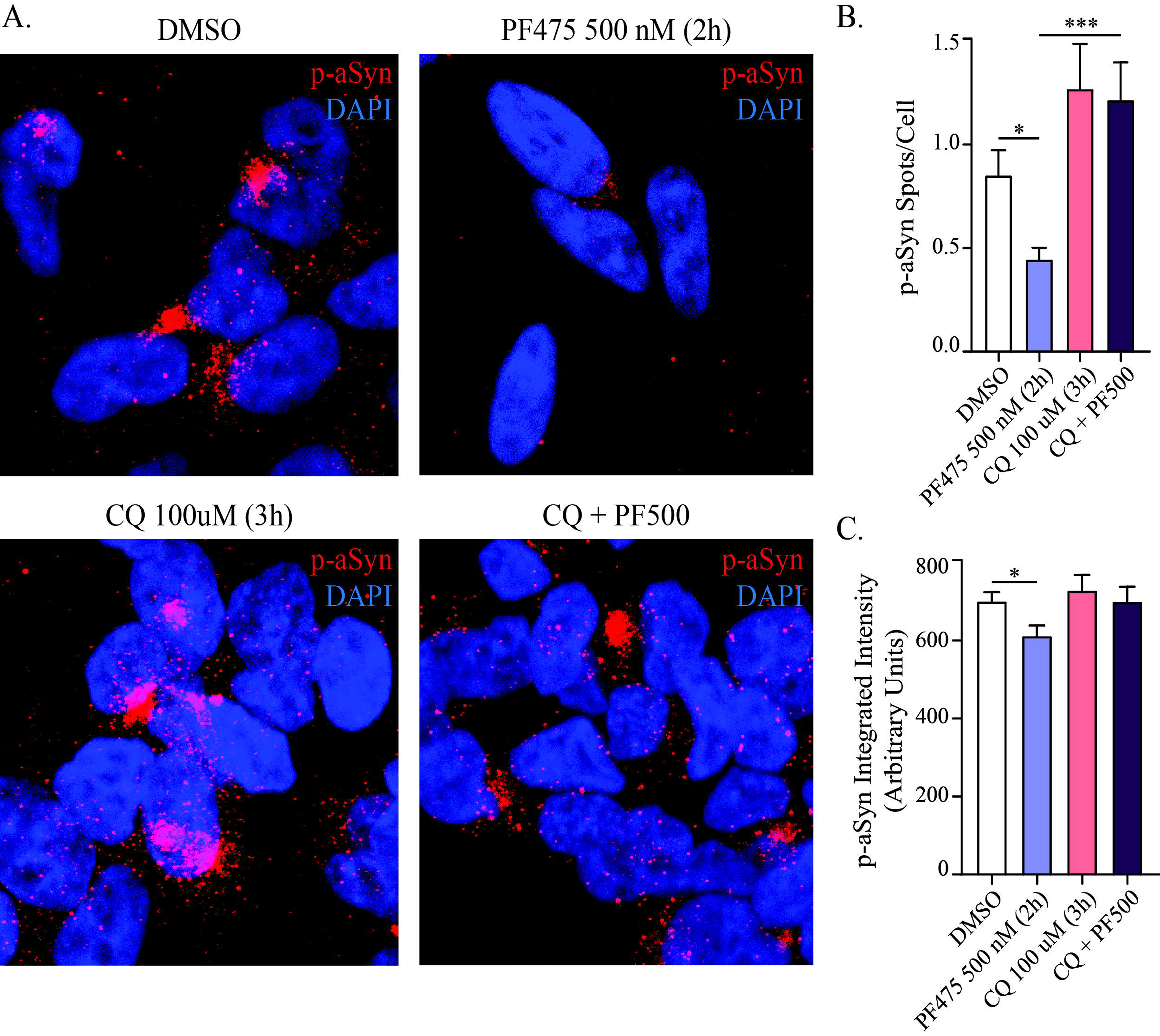
Blockade of autophagosome-lysosome fusion with CQ prevents PF-475-dependent reduction of pS129-aSyn inclusions. A) G2019S-LRRK2 cells were either treated with PF-475 (500 nM, 2h), CQ (100 μM, 3h) or their combination, before processing for pS129-aSyn immunostaining and confocal microscopy. B) The application of PF-475 reduced the number of pS129-aSyn inclusions, while CQ alone did not elicit significant effects. However, when CQ was applied before kinase inhibition, PF-475 lost its efficacy in reducing intracellular inclusion number. C) Similarly, PF-475 alone confirmed its ability in reducing pS129-aSyn signal intensity, while pre-treatment with CQ prevented this effect. Data are means±SEM of 4-5 independent experiments and analysis conducted on 700-1000 cells per group in each experiment. *p<0.05, ***p<0.001, one-way ANOVA followed by Bonferroni’s *post-hoc* test.

This experiment demonstrates that PF-475 reduces aSyn inclusions by acting on the fusion step of the autophagy process, with its upstream blockade completely preventing the positive effect of LRRK2 kinase inhibition.

## Discussion

Our recombinant LRRK2 neuroblastoma cells display an array of abnormalities in the autophagy-lysosome pathway that ultimately impact on the formation of autolysosomes, proteolytic activity and clearance of endogenous pS129-aSyn. The results reported here indicate that G2019S-LRRK2 exerts the more severe effects, causing accumulation of autophagosomes, impairment of lysosome function and accumulation of intracellular pS129-aSyn-positive inclusions. Enhanced cellular levels of WT-LRRK2 also have consequences on this pathway, however cells appear to be capable of better compensation. In fact, despite similar lysosomal depletion, WT-LRRK2 cells retain a strong proteolytic activity, display an enhancement of the autophagosome-lysosome fusion capacity and no detectable pS129-aSyn inclusions are observed. This indicates that the correct advancement of the autophagic process is capable of compensating the lysosomal depletion. Especially, we hypothesize that WT-LRRK2 is not affecting the degradative capacity of remaining lysosomes, as the cells retain the capability of effectively forming autolysosomes. Their functionality is sufficient to carry their task to completion. Consistently, WT- and G2019S-LRRK2 cells display distinct changes in mRNA levels of an array of genes involved in the autophagy-lysosome system. Thus, it can be speculated that enhanced LRRK2 levels *per se* induce variations in this pathway (as LRRK2 likely possesses a role in it), and the G2019S mutation leads to alterations to LRRK2 functionality that reflect in specific functional changes.

It is well established that LRRK2 plays a prominent role in autophagy, endosomal and lysosomal systems. Despite intense research efforts, a definite mechanistic model is still missing, and it is difficult to conclude which parts of the pathway are affected. In contrast to our results, G2019S-LRRK2 has been shown to cause an exaggerated activity of basal autophagy with an increase in autolysosome formation in patient-derived fibroblasts. However, this was not linked to a protective mechanism but rather related to an increase in cytotoxicity. Nevertheless, we report consistent findings regarding increased LC3B-II levels (i.e. increase in autophagosome number) and lysosome defects^20^. Thus, it could well be that the different cellular models employed explain this discordance. Notably, our gene expression array shows an enhancement of *MAP1LC3A* and *MAP1LC3B* transcription in LRRK2 cells, which could be interpreted as an increased autophagic function. However, more targeted functional assays demonstrated the opposite, calling for attention when interpreting results from a single experimental paradigm^36^.

On a similar note, several studies focused on a specific step of the autophagic machinery. Overlap in LRRK2, ERK1/2 and mTOR signaling pathways served as rationale to investigate the role of LRRK2 in autophagy initiation and autophagosome formation and assess the importance of kinase activity^12–14^. Results were contrasting and at the moment it is not possible to draw a definitive conclusion on whether (WT) LRRK2 kinase activity represses or enhances autophagy initiation. As mentioned above, differences in cell models utilized could well underlie this discordance and it is plausible that a cell type-specific regulation exists (e.g. neuron vs astrocyte vs fibroblast).

Our study is not directed towards resolving these issues, but we focused on elucidating the effects of PD-linked G2019S-LRRK2 on the autophagy-lysosome pathway with the specific goal of linking them directly to aSyn clearance.

LRRK2 modulates aSyn neuropathology in a kinase-dependent manner and PD mutations worsen or sensitize to aSyn toxicity^18,26,27,31^. On the other hand, autophagy impairment can play a causative role in aSyn accumulation and nigral neurodegeneration^41–44^. In addition, alterations in autophagy-lysosome markers are found in PD brain areas affected by Lewy pathology^45,46^. However, no studies to date report an experimental demonstration that these phenomena are directly linked. Here, we show that not only does G2019S-LRRK2 cause defects in the autophagy-lysosome pathway in parallel to aSyn accumulation, but we also demonstrate that the formation of autolysosomes is the specific step that is affected. In addition, we provide evidence that these defects are mediated by the (over)activity of LRRK2 caused by the G2019S mutation and its pharmacological inhibition attenuates all the observed abnormalities. Most importantly, these effects are accompanied by reduction of aSyn inclusions.

*Per se*, these results hint that reduction in lysosome function and formation of autolysosomes are causative of aSyn accumulation, but do not yet provide an unequivocal mechanistic demonstration. Thus, we took advantage of the recent clarification that CQ specifically inhibits the fusion between autophagosomes and lysosomes^32^ to further clarify the cellular mechanism in LRRK2 cells. Consistently, upstream blockade of this step completely prevents the reduction in aSyn inclusions operated by the LRRK2 kinase inhibitor PF-475, unequivocally demonstrating that the fusion step is the target of LRRK2 kinase activity and is required for clearance of pathologic aSyn.

Interestingly, other neurodegenerative diseases have been linked to autophagy-lysosome dysfunction mediating accumulation of toxic protein aggregates^47–49^. Specifically, mutant Huntingtin has been reported to also inhibit autophagosome-lysosome fusion in Huntington’s disease^50^, opening fascinating new scenarios of common pathogenic mechanisms across different diseases.

In our study we specifically focused on macroautophagy processing as, in our cell models, it revealed to be the major affected mechanism, with consistent responses to kinase inhibition. Nevertheless, CMA has also been profoundly implicated in LRRK2 biology and aSyn pathology^17,18,41^. Our results do not exclude an involvement of CMA and the possibility that parallel pathways might lead to similar cellular consequences. This hypothesis finds further support from the notion that (macro)autophagy and CMA are functionally related and variations in one cause compensatory change in the other^51,52^.

Lastly, LRRK2 has direct roles on lysosome biology and function^15^ that could “bypass” upstream steps of macroautophagy (consistent with a role in CMA). However, we do observe alterations in WT-LRRK2 cells with regard to lysosome number, but no accumulation of undegraded autophagosomes and/or aSyn inclusions. This could be explained by the ability of forming autolysosomes and the strong proteolytic activity retained by these cells, which could serve as a compensatory mechanism to contrast abnormally elevated LRRK2 levels. We could also speculate that CMA might compensate in WT-LRRK2 cells, retaining an efficacious lysosome functionality that is parallel to autolysosome formation. Thus, a dynamic balance between these pathways could represent an additional (or even synergistic) factor in pathogenesis. Of note, our WT-LRRK2 cells express higher levels of the protein when compared to G2019S-LRRK2 cells^29^, thus it is plausible to expect that cellular processes might be altered. However, kinase activity is overactive in mutant cells despite the difference in total protein content, indicating that cellular processes are more profoundly affected by aberrant enzyme function.

LRRK2 kinase inhibitors have been developed as a disease-modifying therapeutic strategy based on the etiological involvement of increased kinase activity in *Lrrk2* PD patients^38^. Preclinical models confirmed their potential in rescuing toxic effects of mutant LRRK2^53^, providing rationale for clinical trials in familial *Lrrk2* PD. However, recent evidence demonstrated that LRRK2 silencing or kinase inhibition are also effective against aSyn neuropathology and toxicity^26,27,31,54^. Importantly, endogenous LRRK2 has been found overactive in iPD and non-LRRK2 animal models^55^ extending their potential application to all PD patients. On these premises, a clinical trial is ongoing to evaluate a LRRK2 kinase inhibitor in PD patients with and without LRRK2 mutations (ClinicalTrials.gov identifier: NCT03710707). Thus, it is of paramount importance to understand how the inhibitors work from the cellular and molecular mechanistic points of view, also in consideration of their peripheral side effects^53^.

Future work will be directed towards distinguishing the role of endogenous LRRK2 and the effect of point mutations on these pathways, utilizing the evidence presented here to direct research efforts with increased efficacy.

## Acknowledgements

The authors would like to thank Dr. Alexandros Lavdas for the analysis of Lysotracker images using Imaris software. We are also thankful to Anna Masato and Prof. Elisa Greggio (University of Padova) for technical support to Western blotting of pLRRK2.

The authors are also grateful to the Autonomous Province of Bolzano for covering the costs for the Open Access option.

---

**Supp. Fig. 1**. *Autophosphorylation at S935 is not enhanced in G2019S-LRRK2 cells*. A) Western blot analysis was performed to assess phosphorylation levels at S935-LRRK2 in SH-SY5Y, WT-LRRK2 and G2019S-LRRK2 cells. B) The pS935-LRRK2/LRRK2 ratio was significantly lower in G2019S-LRRK2 cells, when compared to WT-LRRK2. Phosphorylation levels of endogenous LRRK2 in control SH-SY5Y cells were too low to yield accurate quantification through band optical density.

Data are means±SEM of 4-5 independent experiments.

*p<0.05, unpaired two-tailed Student’s t-test.

**Supp. Fig. 2**. *LRRK2 kinase inhibition for 6h retains its efficacy in reducing pS129-aSyn inclusions in G2019S-LRRK2 cells*. A) G2019S-LRRK2 cells were treated with PF-475 (300 nM and 500 nM) for 6h, then processed for immunocytochemistry for pS129-aSyn. B) The number of pS129-aSyn inclusion spots is significantly lowered by application of both concentrations of PF-475. C) Similarly, the integrated intensity of the immunosignal is reduced by treatment with PF-475.

Data are means±SEM of 4-5 independent experiments and analysis conducted on 700-1000 cells per group in each experiment.

*p<0.05, **p<0.01, one-way ANOVA followed by Bonferroni’s *post-hoc* test.

