## Supplementary figures and images for "Kinase inhibition of G2019S-LRRK2 restores autolysosome formation and function to reduce endogenous alpha-synuclein intracellular inclusions"

### Supplemental Figure 1

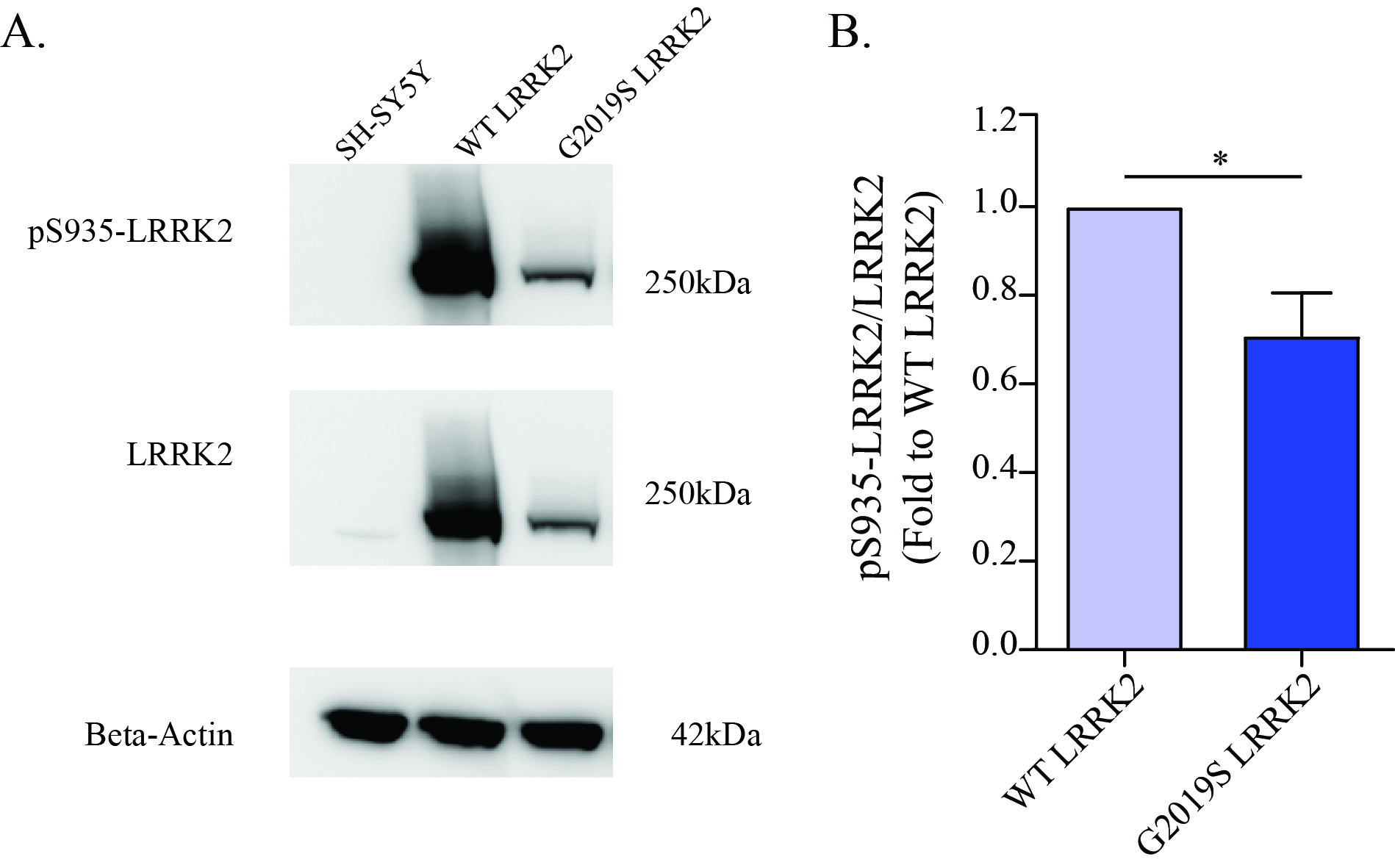

### Supplemental Figure 2

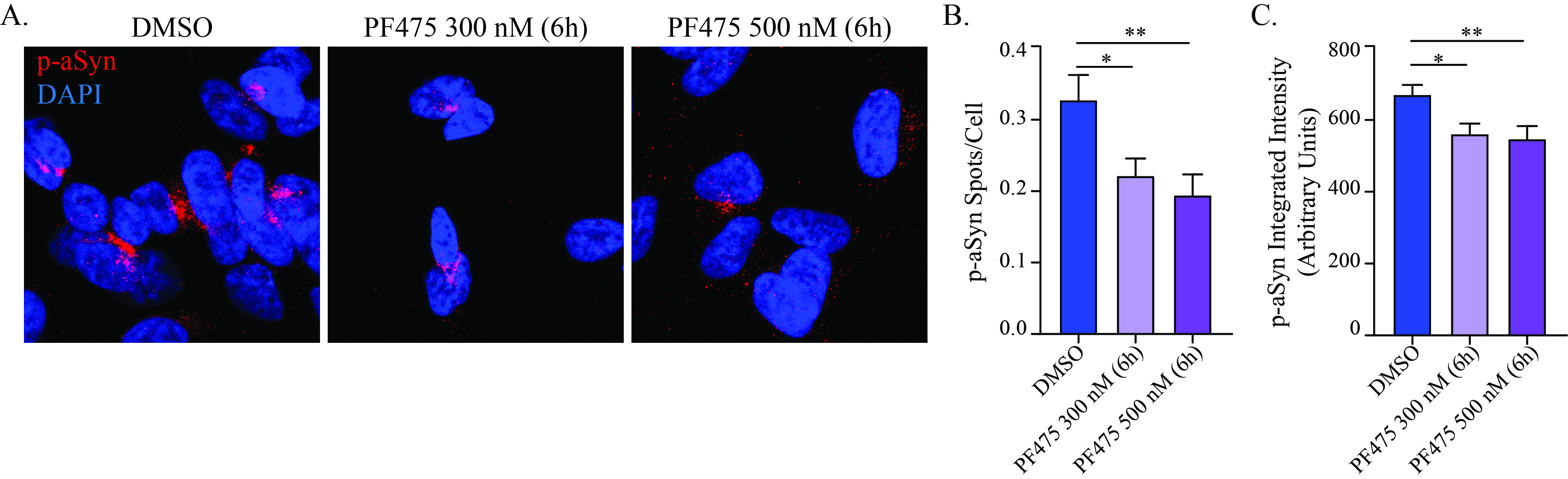
