## Supplemental Table 1 for "Kinase inhibition of G2019S-LRRK2 restores autolysosome formation and function to reduce endogenous alpha-synuclein intracellular inclusions"

| **Symbol** | **Fold Change (comparing to G2019S LRRK2 )** | |
| --- | --- | --- |
|  | **300nM PF475** | **500nM PF475** |
|  | **Fold Change** | **Fold Change** |
| AKT1 | 0.8524 | 0.9129 |
| AMBRA1 | 0.8792 | 0.9659 |
| APP | 0.8552 | 0.8675 |
| ATG10 | 0.8035 | 0.8402 |
| ATG12 | 1.0846 | 1.0418 |
| ATG16L1 | 1.4014 | 1.1839 |
| ATG16L2 | 0.8771 | 1.0325 |
| ATG3 | 0.9923 | 0.9905 |
| ATG4A | 0.8755 | 0.8956 |
| ATG4B | 0.9648 | 0.9781 |
| ATG4C | 0.8971 | 0.8644 |
| ATG4D | 0.9407 | 0.9887 |
| ATG5 | 0.9514 | 0.9305 |
| ATG7 | 0.9705 | 0.9763 |
| ATG9A | 0.9659 | 0.9356 |
| ATG9B | 0.7956 | 0.7643 |
| BAD | 0.8393 | 0.8791 |
| BAK1 | 1.0566 | 1.1624 |
| BAX | 0.8552 | 0.8895 |
| BCL2 | 1.0177 | 1.1083 |
| BCL2L1 | 0.9105 | 0.8696 |
| BECN1 | 0.8571 | 0.8758 |
| BID | 0.948 | 0.988 |
| BNIP3 | 0.9184 | 0.9668 |
| CASP3 | 0.9951 | 1.0347 |
| CDKN1B | 0.9108 | 0.9413 |
| CDKN2A | 1.012 | 0.9644 |
| CLN3 | 0.9608 | 0.9694 |
| CTSB | 1.0006 | 1.0475 |
| CTSD | 0.869 | 0.8911 |
| CTSS | 0.8576 | 0.829 |
| CXCR4 | 0.8962 | 0.8939 |
| DAPK1 | 0.6781 | 0.692 |
| DRAM1 | 0.977 | 0.8721 |
| DRAM2 | 1.0271 | 1.0555 |
| EIF2AK3 | 1.0229 | 1.0854 |
| EIF4G1 | 0.77 | 0.8515 |
| ESR1 | 0.9624 | 0.8185 |
| FADD | 0.7884 | 0.7817 |
| FAS | 0.5322 | 0.592 |
| GAA | 0.7939 | 0.8369 |
| GABARAP | 0.9299 | 0.9443 |
| GABARAPL1 | 1.0293 | 1.0129 |
| GABARAPL2 | 0.9929 | 0.9533 |
| HDAC1 | 0.9149 | 0.9756 |
| HDAC6 | 0.973 | 1.0053 |
| HGS | 0.9496 | 0.8612 |
| HSP90AA1 | 0.9296 | 0.9706 |
| HSPA8 | 1.1158 | 1.0412 |
| HTT | 0.9207 | 0.9367 |
| INS | 2.793 | 0.8725 |
| LAMP1 | 0.871 | 0.8504 |
| MAP1LC3A | 0.8755 | 0.8931 |
| MAP1LC3B | 1.0891 | 1.1389 |
| MAPK14 | 0.9577 | 0.9644 |
| MAPK8 | 1.0091 | 1.0164 |
| MTOR | 0.9361 | 1.0255 |
| NFKB1 | 0.9608 | 0.9907 |
| NPC1 | 0.7853 | 0.8576 |
| PIK3C3 | 0.8869 | 0.8633 |
| PIK3R4 | 0.8912 | 0.8952 |
| PRKAA1 | 0.8933 | 0.9265 |
| PTEN | 0.8542 | 0.8297 |
| RAB24 | 0.7723 | 0.9302 |
| RB1 | 0.8786 | 0.9489 |
| RGS19 | 0.9054 | 0.7684 |
| RPS6KB1 | 0.8521 | 0.9193 |
| SNCA | 0.9224 | 0.8975 |
| SQSTM1 | 1.0964 | 1.2076 |
| TGFB1 | 0.7778 | 0.86 |
| TGM2 | 0.8506 | 0.6638 |
| TMEM74 | 0.7667 | 0.8725 |
| TNFSF10 | 0.8292 | 1.4314 |
| TP53 | 0.8373 | 0.8615 |
| ULK1 | 1.0669 | 1.0377 |
| ULK2 | 0.9798 | 0.9838 |
| UVRAG | 0.8273 | 0.8089 |
| WIPI1 | 0.8926 | 1.0505 |

*Supp. Table 1. Relative expression changes of genes related to the autophagy-lysosome pathway in G2019S-LRRK2 cells treated with 300 nM or 500 nM PF-475, compared to DMSO vehicle control.*
